# Hand preference and the corpus callosum: Is there really no association?

**DOI:** 10.1101/2022.11.14.516402

**Authors:** Nora Raaf, René Westerhausen

## Abstract

Originating from a series of morphometric studies conducted in the 1980s, it appears a widely held belief in cognitive neuroscience that the corpus callosum is larger in non-right handers than in right handers (RH). However, a recent meta-analysis challenges this belief by not finding significant differences in corpus callosum size between handedness groups. Yet, relying on the available published data, the meta-analysis was not able to account for a series of factors potential influencing its outcome, such as confounding effects of brain size differences and a restricted spatial resolution of previous callosal segmentation strategies. To address these remaining questions, we here analysed the midsagittal corpus callosum of N = 1057 participants from the Human Connectome Project (HCP 1200 Young Adults) to compare handedness groups based on consistency (e.g., consistent RH vs. mixed handers, MH) and direction of hand preference (e.g., RH vs. left handers). A possible relevance of brain-size differences was addressed by analysing callosal variability by both using forebrain volume (FBV) as covariate and utilising relative area (callosal area/thickness divided by FBV) as dependent variable. Callosal thickness was analysed at 100 measuring points along the structure to achieve high spatial resolution to detect subregional effects. However, neither of the conducted analyses was able to find significant handedness-related differences in callosal and the respective effect-sizes estimates were small. For example, comparing MH and consistent RH, the effect sizes for difference in callosal area were below a Cohen’s *d* = 0.1 (irrespective of how FBV was included), and narrow confidence intervals allowed to exclude effects above |*d*| = 0.2. Analysing thickness, effect sizes were below *d* = 0.2 with confidence intervals not extending above |*d*| = 0.3. In this, the possible range of population effect sizes of hand preference on callosal morphology appears well below the effects commonly reported for factors like age, sex, or brain size. Effects on cognition or behaviour accordingly can be considered small, questioning the common practise to attribute performance differences between handedness groups to differences in callosal architecture.

## 1 Introduction

Hemispheric specialization is a central feature of human brain organization and individuals differ in the pattern the various cognitive functions are distributed between the hemispheres (Bryden, 1990; Vingerhoets, 2019). The most salient example for this diversity in lateralized brain organization is handedness. While right-hand preference – left hemispheric dominance for the control of skilled manual actions – is prevalent in ca. 90% of individuals, ca. 10% of the population prefer the left hand or do not have a preferred hand (Papadatou-Pastou et al., 2020). In this regard, handedness is special, as it is a form of lateralisation that can be easily assessed, and even in its “atypical” phenotype appears frequently in the population. Thus, handedness is an excellent construct to study neuronal underpinnings of differences in hemispheric lateralization (Willems, der Haegen, Fisher, & Francks, 2014) and a large number of neuroimaging studies identified neuroanatomical asymmetries of grey and whitematter that are linked to handedness (e.g., Amunts, Jäncke, Mohlberg, Steinmetz, & Zilles, 2000; Guadalupe et al., 2014; Hervé, Crivello, Perchey, Mazoyer, & Tzourio-Mazoyer, 2006; Howells et al., 2018; Marie et al., 2015; Steinmetz, Volkmann, Jäncke, & Freund, 1991).

The most studied brain structure in relation to handedness is the corpus callosum (for recent overviews see (Budisavljevic, Castiello, & Begliomini, 2021; Westerhausen & Papadatou-Pastou, 2022)) which, as the major connection between the hemispheres (Schmahmann & Pandya, 2006), is thought to have a central role in establishing (Galaburda, Rosen, & Sherman, 1990; Witelson & Nowakowski, 1991) and supporting hemispheric lateralisation (e.g., Chechlacz, Humphreys, Sotiropoulos, Kennard, & Cazzoli, 2015; Labache et al., 2020; Westerhausen, Gruner, Specht, & Hugdahl, 2009). This focus on corpus callosum morphology originates from a series of influential publications by Sandra Witelson (Witelson, 1985, 1989; Witelson & Goldsmith, 1991). Comparing the corpus callosum of individuals with consistent right-handed (cRH) preference with those of less consistent preference (mixed-handed, MH), she found that MH had a larger corpus callosum than cRH. Witelson’s publications are to date very influential, and studies frequently cite her findings when trying to explain behavioural or cognitive differences between handedness groups (e.g., Jasper, Christman, & Clarkson, 2021; Lyle, Hanaver-Torrez, Hackländer, & Edlin, 2012; Prichard, Propper, & Christman, 2013; Propper, Christman, & Phaneuf, 2005; Roberts, Fernandes, & MacLeod, 2020; Zapała et al., 2020). Replication attempts were, however, rarely successful (Habib et al., 1991), discovered difference in callosal subregions rather than in the total corpus callosum (e.g., Cowell & Gurd, 2018; Jäncke, Staiger, Schlaug, Huang, & Steinmetz, 1997; Luders et al., 2010), or only in one of the two sexes (e.g., Clarke & Zaidel, 1994; Denenberg, Kertesz, & Cowell, 1991). Thus, reviews of the relevant literature typically emphasize the heterogeneity of the findings and question the existence of a general association of callosal architecture and hand preference (Beaton, 1997; Budisavljevic et al., 2021).

Given the above, it appears necessary to discuss reasons why handedness should be associated with structural differences in the corpus callosum. For this purpose, it is important to note that handedness can be conceptualised both as preference for one hand over the other or based on skill (dexterity) differences when performing a manual task (Andersen & Siebner, 2018). Most of the relevant literature, assesses handedness as hand preferences, using self-report questionnaires (as suggested by Annett, 1970; Oldfield, 1971). These questionnaires require to indicate the preferred hand for a range of common manual tasks, such as writing, throwing, using scissors, together with an indication of the consistency of this preference (e.g., “always” vs. “mostly”). The answer pattern can then be summarized as *laterality quotient* (LQ), which expresses the difference in the mount of “left” and “right” answers across the items both in direction (which hand is on average preferred) and strength (consistency of preference across tasks). The use of *direction* of hand preference to classify hand preference appears most intuitive as individuals readily classify themselves referring the preferred hand as left handers (hereafter referred to as dominant left hander, dLH) or right handers (dRH). However, research into handedness is often interested in the strength of the lateralisation irrespective of the direction, referring to the *consistency* of hand preference (e.g., Jasper et al., 2021; Lyle et al., 2012; McDowell, Felton, Vazquez, & Chiarello, 2016). Strong or consistent preference for the left (cLH) or right hand (cRH) across multiple manual tasks is contrasted with inconsistent preference within (ambidexterity) or across tasks (i.e., MH). Inspired by the work of Annett (1970), Witelson (1985, 1989) analysed the corpus callosum based on consistency, comparing MH and cRH participants (and others followed her example, e.g., Habib et al., 1991; Jäncke et al., 1997; Kertesz, Polk, Howell, & Black, 1987; Preuss et al., 2002; Steinmetz et al., 1992)). This comparison allows for a straightforward prediction about the relationship between hand preference and callosal connectivity (Witelson & Nowakowski, 1991). That is, that MH demands more coordination between the hemisphere than consistent hand preference, due to the variation in which hemisphere controls manual performance. Hence, a stronger callosal connection in MH as opposed to cRH (and cLH) would be predicted.

Studies analysing the direction of hand preference in relation to callosal morphology can also be found in the literature (e.g., Denenberg et al., 1991; Luders et al., 2003; Moffat, Hampson, & Lee, 1998; Morton & Rafto, 2006; Nasrallah et al., 1986; Ozdikici, 2020; Westerhausen et al., 2004). Here, however, the theoretical relationship to the corpus callosum is less clear than for consistency approaches. Considering the simplest case, one might assume that the functional organization of the brain of a left- and right-handed individual is in principle identical, just mirrored across the brain’s midline. In this case, the nature of callosal interaction between the hemispheres should be comparable for both hand preference groups so that no structural difference would be predicted. Yet, considering the handedness in interaction with other lateralized functions, it gets obvious that a “fully mirrored” brain organization of dRH and dLH is the exception rather than the rule (Gerrits, Verhelst, & Vingerhoets, 2020; Karlsson, Johnstone, & Carey, 2022). For example, similarly to what is found in dRH, language modules also in dLH are in the majority of cases located in the left hemisphere (although with a substantially reduced prevalence compared to dRH; see (Carey & Johnstone, 2014)). Considering the interaction between language and motor modules (e.g., when writing with the dominant hand), dLH compared with dRH individuals would (viewed on group level) require additional communication between the hemispheres, potentially reflected in stronger structural callosal connectivity. Thus, while the consistency approach predicts corpus callosum morphology to be directly related to hand-motor functioning, the direction approach requires assumptions about the interaction with additional functional modules to come to a comparable prediction.

Reflecting the general heterogeneity of findings in the literature (Beaton, 1997; Budisavljevic et al., 2021), a recent meta-analysis did neither for consistency-nor direction-based hand preference groups find statistical significant differences in corpus callosum size (Westerhausen & Papadatou-Pastou, 2022). That is, considering consistency comparing cRH and MH samples the estimated effect size, expressed as Hedges’ *g*, −0.004 in favour of MH, while for direction comparison of dRH and dLH the estimate was g = 0.02. The confidence intervals of the mean estimates suggested that in both cases population-effect sizes below g = −0.20 and above 0.20 can be excluded with reasonable confidence. Thus, interpreting these effect sizes to be small and likely irrelevant, one feels tempted to conclude that hand preference and callosal morphology are not associated to a theoretically relevant degree. However, the meta-analysis also identified a series of factors that might alter this conclusion, but which could not be addressed by the meta-analysis as relevant data was not available in the primary literature. Firstly, the meta-analysis integrated studies using absolute size (i.e., absolute midsagittal area or volume measures) of the corpus callosum as dependent variable, while a possible moderating or confounding influence of brain-size variability (e.g., Jäncke, Mérillat, Liem, & Hänggi, 2015; Jäncke et al., 1997; Witelson, 1989) could not be evaluated. Secondly, considering selective subregional (as opposed to the entire corpus callosum) associations, the meta-analysis was bound to the callosal parcellation schemas used in the literature so that the spatial sensitivity was not sufficient to evaluate small regional effects that do not adhere to the predefined parcellation schemas (for an example see Luders et al., 2010).

The present study was conceptualised as a follow-up of the discussed meta-analysis to evaluate the open questions in a series of interrelated analyses of the association of hand preference and corpus callosum measures using data from the Young Adult sample of the Human Connectome Project (HCP, Van Essen et al., 2013). Hand preference, as independent variable, was determined based on the assessment with the HCP’s version of the Edinburgh Inventory (EHI, Oldfield, 1971). Hand preference groups were defined based on consistency (i.e., cRH vs. MH, using two competing definition of hand-preference consistency) and direction of hand preference (dRH vs. dLH, and cRH vs. consistent left handers, cLH) following the most common approaches in the literature. To comprehensively address possible confounding or moderating effects of brain size (i.e., open question 1), the corpus callosum was assessed both using forebrain volume (FBV) as covariate in the analysis of its absolute midsagittal area, as well as by using relative area (absolute area divided by FBV) as dependent variable. To examine regional differences with good spatial resolution (question 2), cortical thickness was analysed at 100 measuring points, by analysing absolute callosal thickness measures with and without FBV as covariate, along with thickness to FBV ratios.

## 2 Methods

### 2.1 Data and participants

The sample consists of the 1113 of 1206 participants of the Young Adult HCP sample (Van Essen et al., 2013) for which structural MRI data was available. Of these, 47 (4.2%) were excluded due to known anatomical anomalies (see https://wiki.humanconnectome.org/), and 9 (0.8%) as the corpus callosum extraction was not successful. This leaves a final sample of *N* = 1057 participants (581 females, 476 males) with a mean age of 28.8 years (standard deviation, *sd* = 3.7 years; range: 22 to 37 years).

### 2.2 Hand preference assessment

The HCP assessed hand preference using a version of the EHI (Oldfield, 1971) which differed from the original version in the number of items and response format. The number of items was reduced from ten to nine, leaving out the item asking for “drawing” (see also (Ruck & Schoenemann, 2021)). The response format was the commonly used 5-scale answer format (adopted from (Schachter, Ransil, & Geschwind, 1987)) with the answer options “always left”, “usually left”, “no preference”, “usually right”, and “always right”. Of note, as outlined in Supplement Section A, the variable “Handedness” provided by HCP represents an LQ calculated including the EHI item “kicking with foot” in addition to the nine “manual” items, so that it represents a mixture of manual and pedal preference. Thus, we were forced to calculate a new LQ variable based on the nine manual items, which was achieved by double weighing the answer to the item “writing” (see Supplement Section A for the reasoning behind this choice).

The newly calculated LQ ranged from −100 for strong left-hand preference to 100 for strong right-hand preference. The frequency distribution for the N = 1057 included participants, Figure 1, follows the typical J-shaped pattern: Participants with strong righthand preference dominate the distribution, with few MH, and a slightly elevated number of strong left-handers. The median was accordingly at an LQ of 80, with a mean of 66.7 (sd = 47.2).

**Fig. 1.**
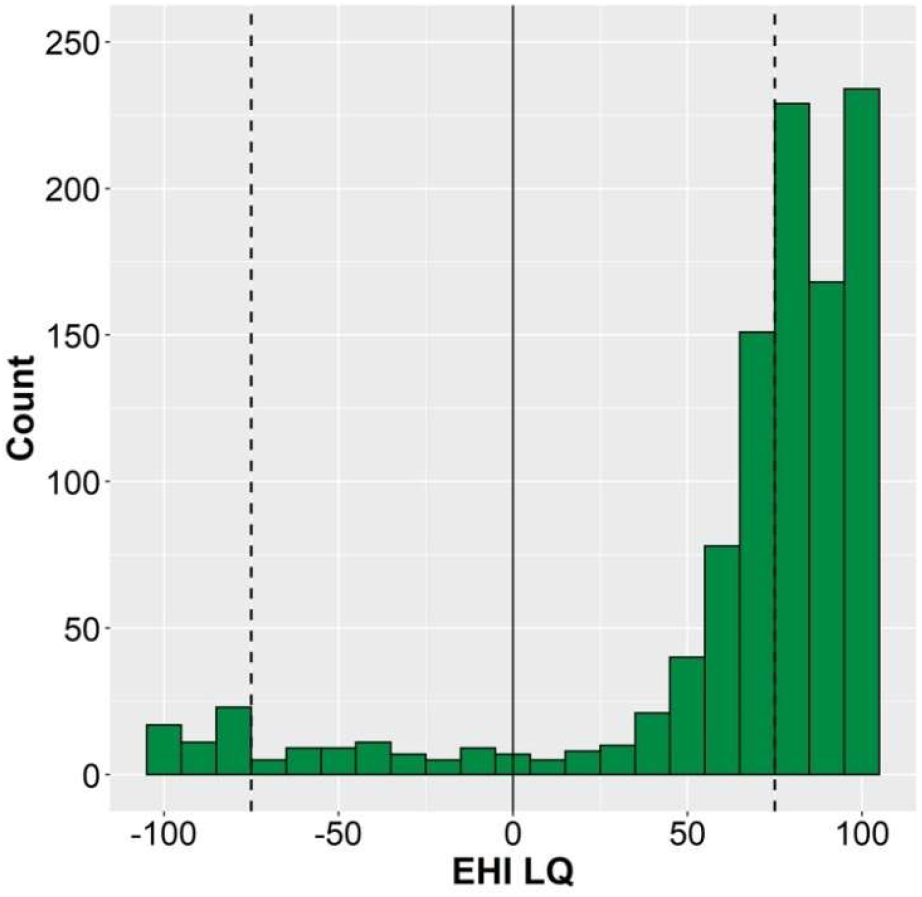
Frequency distribution of the newly calculated EHI LQ (N = 1057). The LQ ranges from −100 (extreme preference for the left hand) to 100 (extreme preference of the right hand). The vertical dashed lines indicate the LQ = 80 cut-off used by Habib et al. (1991) to define consistent left- and right-hand preference, respectively.

### 2.3 Classification of handedness groups

It is common practise in the corpus callosum literature to classify individuals into handedness groups rather than using the LQ as continuous variable. One reason for this can be found in the influential work by Annett (Annett, 1970, 1972), emphasizing the qualitative over gradual difference in hand preference. Also, the J-shaped distribution of the LQ renders an appropriate statistical analysis as continuous predictor difficult (Dragovic & Hammond, 2007). As one main aim of the present project was to replicate previous work, we also here relied on handedness classification. For this purpose, coding variables based both on consistency and direction of hand preference were created as outlined in the following.

Consistency-based coding may either be based on a qualitative analysis of the individual answer pattern (e.g., Clarke & Zaidel, 1994; Jäncke et al., 1997; Witelson, 1989) or be based on applying a specific LQ value as threshold (e.g., Habib et al., 1991; McDowell et al., 2016; Welcome et al., 2009). Witelson (Witelson, 1985, 1989) defined *“… consistentright-hand preference […] as all ‘right’, or ‘right’ with some ‘either’ preferences […] non consistent-right hand preference […]as left-hand preference on at least one of the twelve tasks, regardless of the hand used in writing”* (p. 803, Witelson, 1989). Using the nine items available here, we formed a grouping variable that is comparable to the Witelson classification. That is, all individuals who indicated left-hand preference (“always left” or “usually left”) for at least one task were classified as NcRH. Individuals who had answers on most items with “no preference” (so five or more of the nine items) were also classified as NcRH. The remaining participants were defined as cRH. This led to the classification of 636 participants (or 60.2% of total sample) as cRH and 421 as NcRH. Defining cLH, as NcRH which do not indicate right-hand preference (“always right” or “usually right”) for any of the items, further divided the NcRH sample into 385 MH (36.4% of total sample) and 36 cLH (3.4%) participants.

Defining a LQ cut-off to separate consistent cRH from MH is arbitrary and threshold values between 75 (e.g., Propper & Christman, 2004) and 100 (e.g., Welcome et al., 2009) have been used. In the aim to replicate a study that reported handedness group difference, we here follow Habib et al. (1991) in that *“subjects were first divided into two groups: consistent right-handers, including all subjects having a LQ of+80 or more, and nonconsistent right-handers, including all other subjects.”* (p. 46). For the present data this resulted in 631 participants being classified as cRH (59.7%) and 426 as NcRH. The NcRH group was then further split by classifying all LQ of −80 and below as cLH, yielding 36 cLH (3.4%) and 390 MH (36.9%) participants (see Fig. 1 for illustration of the cut-off in relation to the distribution).

Taken at face value, Habib criterion yields a frequency distribution which is very similar to the distribution obtained using the Witelson criterion. However, the two cannot be used interchangeably as 277 (26.2%) of the participants were classified differently when using the Habib compared to the Witelson criterion (for details Supplement Section B, and Supplementary Table S2). Thus, separate analyses were set up for the two classifications.

The direction of hand preference has mainly been defined by using the preferred hand for writing (e.g., Cowell & Gurd, 2018; Denenberg et al., 1991) or the overall lateral preference across multiple activities (e.g., as expressed by a positive or negative LQ) as indicator (e.g., Martens, Wilson, Chen, Wood, & Reutens, 2013; Moffat et al., 1998; Nasrallah et al., 1986). A supplementary analysis, comparing both approaches produce largely overlapping classifications in the present data (see Supplement Section B, Supplementary Table S3). Consequently, we here use the classification based on the sign of the LQ. That is, 946 (89.5%) participants with a positive LQ were classified as right-dominant (dRH) and 106 (10.0%) with a negative LQ as left-hand dominant (dLH). Five (0.5%) individuals with an LQ of 0 were classified as having “no preference”.

### 2.4 Structural imaging data from HCP

The HCP T1-weighted data obtained on the 3T Siemens Skyra “Connectome” scanner were utilised in the present analyses. Details and considerations behind the applied imaging sequences can be found in HCP reference papers (Glasser et al., 2016; Glasser et al., 2013). In brief, the T1 images were acquired using a magnetization prepared rapid gradient echo (MPRAGE; parameters: repetition time, TR = 2400 ms; echo time, TE = 2.14 ms; inversion time, TI = 1000 ms; flip angle = 8°, iPAT = 2) in 256 sagittal slices (0.7 mm thickness) with field of view of 224 x 224 mm (scan matrix = 320 x 320). Thus, the resolution in the sagittal plane, which is relevant for the assessment of the corpus callosum, was 0.7 x 0.7 mm^2^

### 2.5 Extraction of morphological measures of the corpus callosum

The midsagittal corpus callosum was extracted from the T1-weighted images using the following processing step. Firstly, using the routines provided with the Corpus Callosum Thickness Profile Analysis Pipeline (*CCSegThickness*, version 2.0; (Adamson, Beare, Walterfang, & Seal, 2014)), the midsagittal plane was automatically identified and the corpus callosum extracted. In a second step, the software’s manually editing tool was used to visually inspect and, where necessary, correct the segmentation obtained from the previous step. Misclassifications of pericallosal veins or the fornix, were the most common segmentation errors that required correction. The corrected segmentations where then exported as binary masks in native space. Using routines written in Matlab (version: R2018b; MathWorks Inc. Natick, MA, USA), the number of voxels within this mask multiplied with the voxel-surface area of 0.49 mm^2^ in the sagittal plane was determined for each individual representing the total midsagittal area of the corpus callosum. This procedure has been shown to yield an inter-rater reliability of *r_icc_* =.96 (intra-class correlation calculated as two-way random effects, considering the single measure and absolute agreement; see (Danielsen et al., 2020)). In the present sample, the measured corpus callosum areas ranged from 435.6 to 1002.1 mm^2^ with a mean area of 691.3 (sd: 94.1) mm^2^.

To determine callosal thickness, the outline of each individual corpus callosum was determined removing all non-border voxels. On this outline, the tip of the rostrum (i.e., the posterior-most point of the in-bend anterior half of the corpus callosum) as well as the base of the posterior corpus callosum (i.e., the ventral-most voxel in the splenium) were identified automatically but corrected manually where necessary (Westerhausen et al., 2016; Westerhausen et al., 2018). The tip of the rostrum and base of the splenium then served to divide the callosal outline in a ventral and dorsal part. A midline between the ventral and dorsal outlines was then calculated as mean coordinate of 100 sampling points spaced equally along the two outlines. Finally, regional thickness (i.e., distance between the two outlines) was determined orthogonal to this midline, sampled at 100 equidistantly spaced measurement points. The number of measurement points was selected to be comparable to the spatial resolution used by previous handedness which used 99 or 100 measurement points (Cowell & Gurd, 2018; Denenberg et al., 1991; Luders et al., 2010).

### 2.6 Forebrain volume (FBV)

As measure of brain-size difference we utilised the supratentorial volume as provided with the HCP data, which henceforth is referred to as forebrain volume (FBV). The FBV extraction of the data release was done with the HCP Pipeline (v3.21; for general overview see: (Glasser et al., 2013)) including FreeSurfer 5.3.0. For the present analyses, FBV was preferred over intra-cranial volume or total brain volume, as it mainly includes brain compartments which are connected via callosal fibres, while it excludes those brain structures (i.e., cerebellum and brain stem) which do not have such connections (Schmahmann & Pandya, 2006). Consequently, callosal size is more strongly related with FBV than with intracranial volume (Jäncke et al., 2015). In the present sample, FBV was correlated with area and thickness measures of the corpus callosum explaining 13% of variance of total area and up to 9% of regional thickness measures (see Supplement Section C for details, including Figurs S1 and S2). Thus, an adjustment of the present analysis for potential confounding effects of FBV variability appeared justified. However, callosal variables represent area and distance (thickness) measures, so that FBV (i.e., a volume) had to be converted to allow for an unbiased correction between the different dimensionalities (Jäncke et al., 1997; Smith, 2005). That is, for analysis using callosal area as dependent measure, FBV was raised to the power of 2/3 (i.e., FBV^2/3^), while for thickness analyses the exponent 1/3 (i.e., FBV^1/3^) was used for the conversion.

### 2.7 Statistical analysis

Difference in corpus callosum architecture in relation to hand preference were examined by realising the following four comparisons across the two dependent measures of total callosal area and regional thickness:

- *Comparison A* contrasts cRH and MH individuals, as defined by Witelson’s qualitative criterion. As the original publication (Witelson, 1985, 1989) did not include cLH individuals, they were also excluded here.
- *Comparison B* compares cRH and MH following Habib’s quantitative criterion. This approach was introduced to evaluate the effect of quantitative definition of consistency as compared to the qualitative definition in Comparison A, as well as an attempt to replicate the findings by Habib et al. (1991).
- *Comparison C* focusses on the direction of hand preference by comparing dRH and dLH individuals, as defined using the sign of the EHI for classification. This approach follows a series of previous studies (e.g., Martens et al., 2013; Moffat et al., 1998; Nasrallah et al., 1986).
- *Comparison D* contrasts cRH and cLH individuals according to the Witelson criterion. This approach again follows previous studies (Jäncke et al., 1997; Luders et al., 2003; Steinmetz et al., 1992), and allows to evaluate the effect of direction of hand preference controlling for differences in the consistency which might act as confounding factor in Comparison C (i.e,, the absolute value of the LQ of dLH is lower than in dRH).

The analyses of total corpus callosum area was conducted three times per comparison: twice using absolute area as dependent variable while for one of the two analyses the converted FBV (i.e., FBV^2/3^) was added as covariate, as well as ones using relative area (i.e., area/FBV^2/3^) as dependent variable. Likewise, the segment-wise analyses of regional thickness were conducted for absolute thickness (without and with FBV^1/3^ as covariate) and for relative thickness (i.e., thickness/FBV^1/3^).

The basic design of all analyses was a two-factorial analysis of variance (ANOVA) design with the between factors Handedness Group and Sex. Sex was included as some studies had reported sex differences in the handedness effect on the corpus callosum (Burke & Yeo, 1994; Clarke & Zaidel, 1994; Denenberg et al., 1991; Witelson, 1989). When including the converted FBV as covariate, the analysis was run as analysis of covariance. The participants age was not added as covariate as in the age range of the sample, age effects do not explain substantial variance in the corpus callosum (Danielsen et al., 2020; Skumlien, Sederevicius, Fjell, Walhovd, & Westerhausen, 2018).

All statistical analyses were conducted in R 4.1.0, and scripts of all analyses are available via the accompanying Open Science Forum project (https://osf.io/u25gn/). Segment-wise thickness were corrected for multiple comparisons by adjusting to a False Discovery Rate (FDR) to 5% using the Benjamini-Holberg method, again following a previous study (Luders et al., 2010). As this adjustment is somewhat conservative and we are interested in high statistical sensitivity, we additionally indicated segments/voxels which would be significant if uncorrected p-values would be used as decision criterion. Effect size for Handedness Group and Sex main effects were expressed as Cohen’s *d* based on the estimated marginal means as determined using the *emmeans* package (Lenth, 2021). The Handedenss Group effects are coded so that negative *d*-values reflect larger means in the right-handed groups (cRH, dRH), and positive *d*-values larger means in the non-right handed sample. The 95% confidence intervals (*CI_95%_*) are provided with the point estimates of the effect size to reflect the precision of the estimate (Cumming, 2014). The effect size for interaction effects was expressed as explained variance (*η^2^*).

### 2.8 Data availability statement

The data used in the present analysis can be downloaded in tabulated and deidentified form from the linked Open Science Forum project (https://osf.io/u25gn/). The MRI raw data and other variables used are publicly available via HCP webpage after registration (https://www.humanconnectome.org/).

## 3 Results

### 3.1 Total midsagittal area of the corpus callosum

Considering the consistency-based comparisons A and B, Cohen’s *d* for the main effect on absolute area was −0.04 and −0.06 in favour of cRH over MH for comparison A and B, respectively, with all uncorrected p-values above .38. Test statistics and estimated marginal means are presented in Table 1. For the analyses considering brain size, all *p*-values for the main effect were above .30. The effect size varied between *d* = −0.04 and −0.07 when the converted FBV was added as covariate, and between *d* = −0.04 and −0.06 for relative area. In all cases the estimated marginal means were larger in the cRH than the MH sample.

**Table 1.** Results of the ANOVA’ANCOVA analysing total midsagittal area of the corpus callosum; main effect of Handedness Group (HG) and interaction of HG with Sex are shown.

| DV/COV | Comparison | Estimated Means (SE)<sup>a</sup> |  | Main effect Handedness Group |  |  |  |  | Interaction effect<sup>b</sup> |  |
| --- | --- | --- | --- | --- | --- | --- | --- | --- | --- | --- |
| | | RH group | Other | *F*-value | df<sub>n</sub>; df<sub>d</sub> | *p* | *d* | *CI*<sub>95%</sub>(*d*) | *p* | $\eta^2$ |
| DV: absolute area | A: cRH vs. MH | 694 (3.79) | 690 (4.79) | 0.37 | 1; 1017 | 0.54 | -0.04 | -0.17; 0.09 | 0.98 | <0.001 |
|  | B: cRH vs. MH | 695 (3.88) | 689 (4.83) | 0.77 | 1; 1017 | 0.38 | -0.06 | -0.19; 0.07 | 0.53 | <0.001 |
|  | C: dRH vs. dLH | 693 (3.07) | 681 (9.11) | 1.76 | 1; 1048 | 0.18 | -0.14 | -0.34; 0.07 | 0.77 | <0.001 |
|  | D: cRH vs. cLH | 694 (3.83) | 678 (15.89) | 0.95 | 1; 668 | 0.33 | -0.17 | -0.51; 0.17 | 0.24 | 0.002 |
| DV: absolute area<br>COV: FBV<sup>2/3</sup> | A: cRH vs. MH | 691 (3.52) | 688 (4.44) | 0.42 | 1; 1016 | 0.52 | -0.04 | -0.17; 0.09 | 0.78 | <0.001 |
|  | B: cRH vs. MH | 692 (3.60) | 686 (4.48) | 1.09 | 1; 1016 | 0.30 | -0.07 | -0.20; 0.06 | 0.79 | <0.001 |
|  | C: dRH vs. dLH | 690 (2.84) | 683 (8.41) | 0.66 | 1; 1047 | 0.42 | -0.08 | -0.29; 0.12 | 0.66 | <0.001 |
|  | D: cRH vs. cLH | 687 (3.57) | 681 (14.57) | 0.15 | 1; 667 | 0.70 | -0.07 | -0.41; 0.27 | 0.49 | <0.001 |
| DV: relative area | A: cRH vs. MH | 6.78 (0.03) | 6.75 (0.04) | 0.32 | 1; 1017 | 0.57 | -0.04 | -0.17; 0.09 | 0.79 | <0.001 |
|  | B: cRH vs. MH | 6.79 (0.04) | 6.74 (0.04) | 0.88 | 1; 1017 | 0.35 | -0.06 | -0.19; 0.07 | 0.84 | <0.001 |
|  | C: dRH vs. dLH | 6.78 (0.03) | 6.72 (0.08) | 0.42 | 1; 1048 | 0.52 | -0.07 | -0.27; 0.14 | 0.86 | <0.001 |
|  | D: cRH vs. cLH | 6.78 (0.03) | 6.72 (0.14) | 0.16 | 1; 668 | 0.69 | -0.07 | -0.41; 0.27 | 0.65 | <0.001 |

Notes. (a) estimated marginal means of absolute area is expressed in mm<sup>2</sup>, while the relative area does not have a unit; (b) only $p$ -value and effect size provided here; full statistics can be found in the accompanying documentation together with the test statistics of the main effect of sex (https://osf.io/u25gn/). Abbreviations: DV, dependent variable; COV, covariate; FBV, forebrain volume; df, degrees of freedom

Regarding the direction-based comparisons C and D, the effect size of the main effect of Handedness Group were slightly larger than for the consistency comparisons and varied between *d* = −0.14 and −0.17 for absolute area, suggesting larger area in right- (dRH/cRH) as compared with the left-handed (dLH, cLH) samples. The effect size was *d* = −0.08 and −0.07 when converted FBV was included as covariate, and *d* = −0.07 for both comparisons of relative area. Across all direction comparison the *p*-values of the Handedness Group main effect were above .18. Of note, for comparison D, the overall smaller sample size resulted in low precision of the effect size estimate as reflected in wide confidence intervals, and effects up *d* = −0.51 (for absolute area) cannot be excluded with confidence.

The main effect of sex was significant in all analyses. As the analysis of the Sex effects is repeated in a comparable way in all analyses, the presentation here focusses on the analysis with the largest sample size (analyses regarding comparison C) to avoid redundant reporting. Here, the main effect on absolute area had an *F_1,1048_* = 4.85, was significant (p = .03), and indicated larger overall corpus callosum area in males (estimated marginal mean: 698 mm^2^, ± standard error: 6.66) compared with females (676 ± 6.93 mm^2^). The effect size was *d* = −0.23 (*CI_95%_*: −0.43; −0.02). Introducing FBV as covariate the main effect was significant (*F_1,1047_* = 17.58, *p* < .001) but now indicating larger corpora callosa in females (707 ± 6.80 mm^2^) compared to males (665 ± 6.60 mm^2^) equivalent to an effect size of *d* = 0.49 (*CI_95%_*: 0.26; 0.72). Considering relative area, the main effect of Sex was also found significant (relative area: *F_1,1048_* = 28.01, *p* < .001) with larger corpus callosum to FBV ratios in females (6.98 ± 0.06) than males (6.52 ± 0.06). Here, the effect size was *d* = 0.54 (*CI_95%_*: 0.34; 0.75).

Finally, the interaction of Handedness Group and Sex was not significant in any of the analyses as shown in Table 1. Across all analyses, the interaction effect explained less than 0.2% of the variance (i.e., *η^2^* < 0.002).

### 3.2 Regional callosal thickness measures

Comparable to the area analysis, in neither of the comparisons a significant main effect of Handedness Group was detected on regional thickness. Figures 2 and 3 provide an overview of the effect size per thickness segment, for the analyses focus on consistency and direction of hand preference, respectively.

**Fig. 2.**
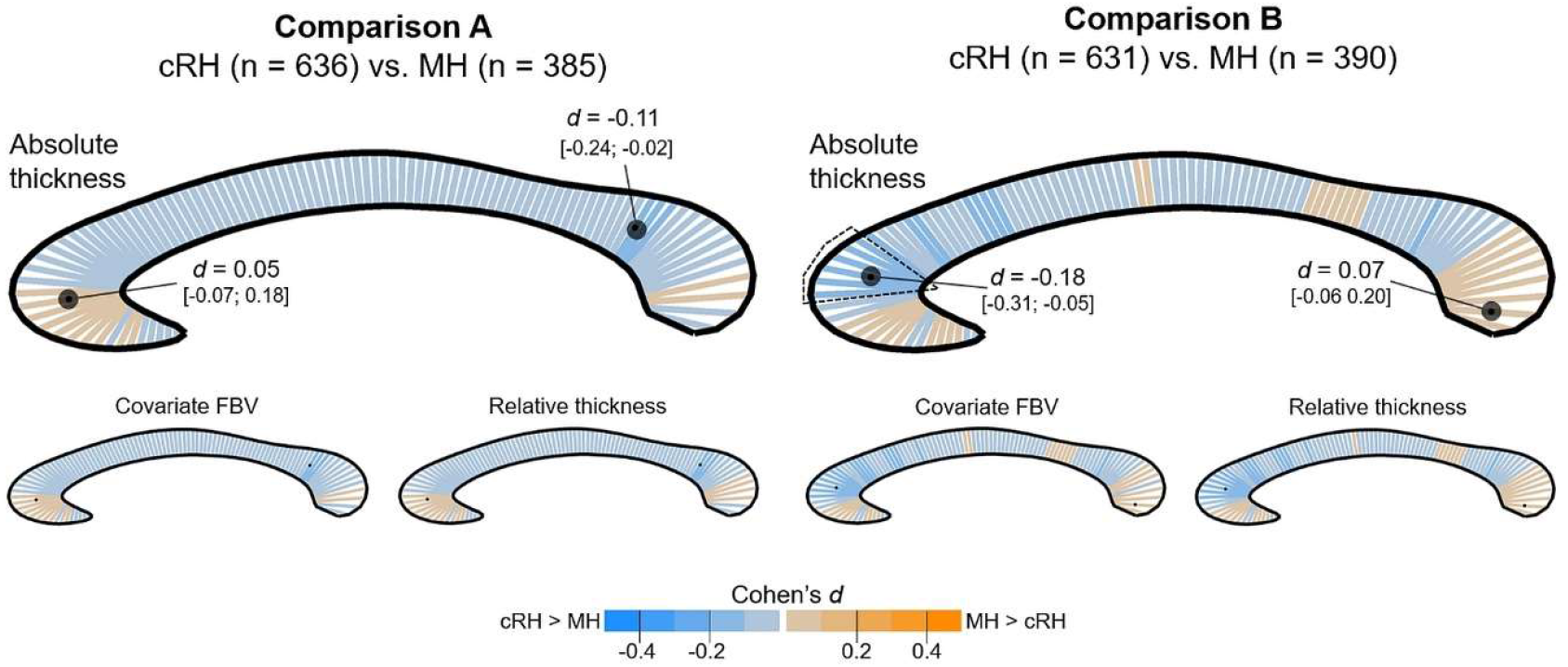
Segment-wise color-coded presentation of the effect size (Cohen’s d) of the Handedness Group main effect for the consistency-based definition of handedness. The colour orange codes differences in favour of the MH group, while blue codes difference in favour of the cRH group. The location of the minimum and maximum effect size are marked by black dots on the respective segment. For the absolute thickness analysis (top), the value of minimum and maximum *d* are additionally provided together with their 95 confidence intervals. Of note, for neither of the analyses did the main effect survive FDR adjustment. For a small number of individual segments, uncorrected p-values were below 0.05 and respective segments are surrounded by a dotted line.

**Fig. 3.**
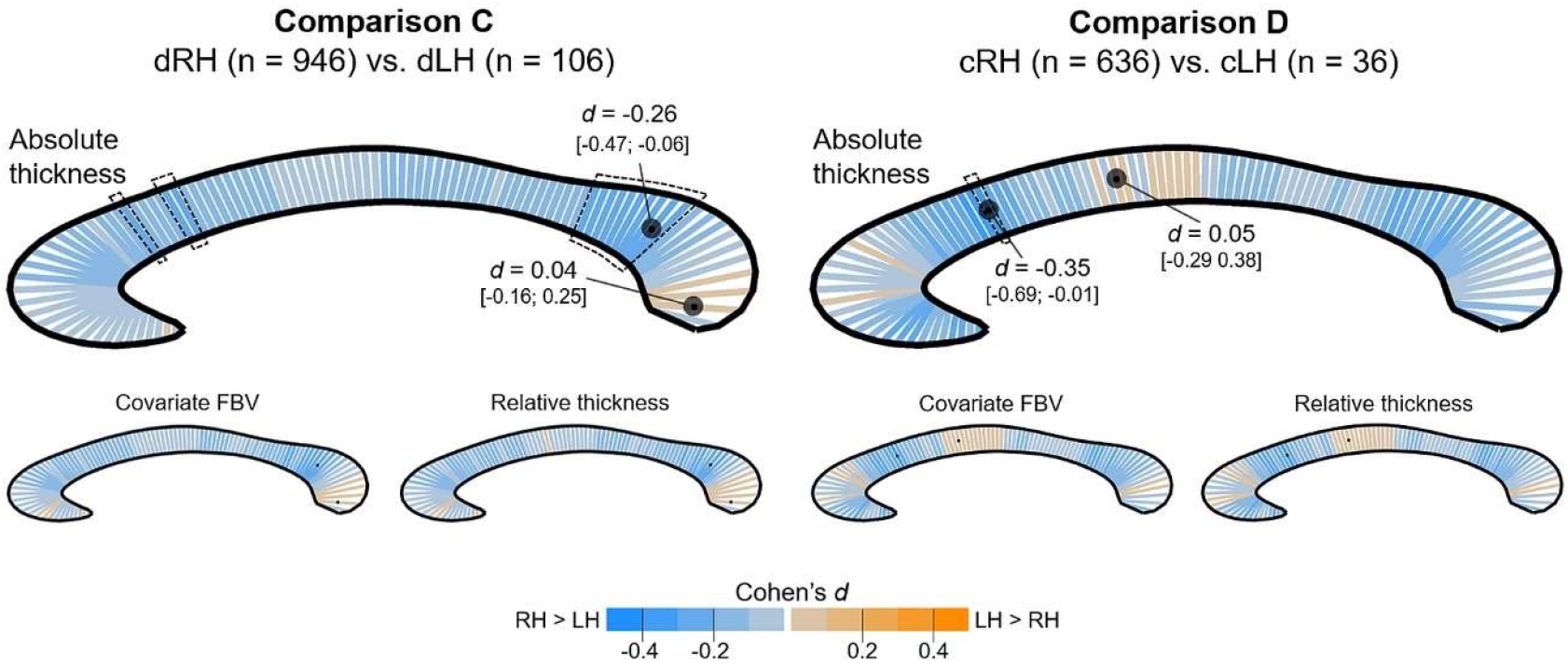
Segment-wise presentation of the effect size of the Handedness Group main effect for the direction-based definition of handedness. Orange indicates mean deviations in favour of the LH group (dLH/cLH), while blue codes difference in favour of the RH group (dRH/cRH). Location of minimum/maximum *d* are marked with black dots. For the absolute thickness analysis (top), the minimum/maximum values (and 95% confidence limits) are additionally provided. For neither of the analyses did the main effects survive FDR adjustment. For some segments the uncorrected p-values were below 0.05. These are surrounded by a dotted line.

Exploring the effect size for the consistency-based comparisons, the estimated Cohen’s *d* in comparison A ranged from *d* = −0.11 (*CI_95%_*: −0.24; −0.02) in favour of cRH (segment location: dorsal splenium) to *d* = 0.05 (*CI_95%_*: −0.07; 0.18) in favour of MH (genu). In comparison B, several segments in the genu were significant before FDR adjustment, whereby the maximum effect size in these segments was *d* = −0.18 (*CI_95%_*: −0.31; −0.05) with increased regional thickness in cRH. The maximum effect size favouring MH (located in the ventral splenium) was *d* = 0.07 (*CI_95%_*: −0.06; 0.20). Considering brain size in these analyses did not substantial change the overall pattern nor the effect size of the findings as can be seen in the small inlay of Fig. 2.

Considering the direction of hand preference (comparisons C and D; Fig. 3), effect sizes were larger than in the consistency-based analyses, with a shift towards thicker callosal segments in the cRH/dRH than the cLH/dLH groups, while remaining non-significant after FDR adjustment. Regarding comparison C, a cluster of segments in the dorsal splenium region were significant before FDR correction, with the maximum effect size being *d* = −0.26 (*CI_95%_*: −0.47; −0.06) in favour of dRH. The maximum effect in the opposite direction was *d* = 0.04 (*CI_95%_*:: −0.16; 0.25). In comparison D, the maximum effect was *d* = −0.35 (*CI_95%_*:: −0.69; −0.01) indicating an increased thickness in cRH than in cLH and was found in a segment located in the dorsal genu. This was also the only segment with an uncorrected *p-*value below 0.05. In the opposite direction, the maximum effect was *d* = 0.05 (*CI_95%_*: −0.29; 0.38). Again, including FBV as covariate or by using relative thickness measures as dependent variable, did not affect the overall pattern of the findings as shown in Fig. 3.

Segments with a significant main effect of Sex (here only presented for the analysis testing comparison C as it has the largest sample) were found for absolute and relative thickness measures, as shown in Fig. 4. Regarding absolute thickness, only three segments in the dorsal splenium survived multiple comparison correction, suggesting a thicker corpus callosum in males compared with females (max effect size *d* = −0.41). Additionally, clusters of segments in genu and ventral splenium were significant before adjustment, also suggesting thicker male corpora callosa.

**Fig. 4.**
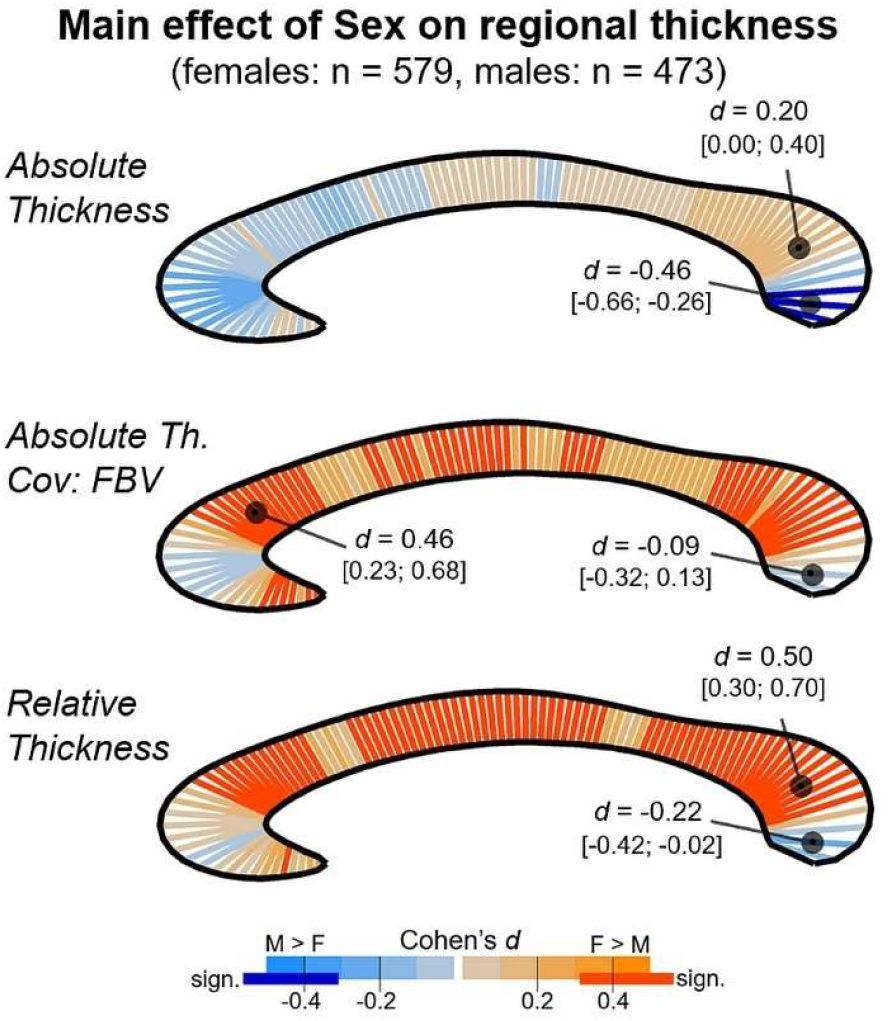
Segment-wise presentation of the effect size *d* of the main effect of Sex for the three analyses conducted in relation to comparison C. Orange codes thicker female, blue thicker male segments. Minimum/maximum *d* values (and 95% confidence limits) are additionally provided). Dark orange and dark blue indicate significant segments after FDR adjustment to 5%.

As can be seen in Fig 4, middle panel, adding FBV^1/3^ as covariate to the analysis changes the overall pattern by shifting the difference from favouring males (coded in blue colour) to favouring females (orange). The sex effect was significant after FDR correction in 54 segments split up in multiple clusters distributed across the entire corpus callosum. The effect size of significant segments ranged from *d* = 0.26 to 0.46. Comparably, the analysis of relative thickness, revealed 68 segments showing a sex difference in favour of females which were distributed in three main clusters located in genu, truncus, and splenium (effect sizes between *d* = 0.22 and 0.50).

For neither of the above analyses, segments revealing a significant interaction of Handedness Group and Sex were detected. The maximum effect found for the interaction effect for any analysis/segment was *η^2^* = 0.005.

## 4 Discussion

The present study examined the effect of handedness on the corpus callosum by empirically addressing two research questions that could not be conclusively answered in a recent metaanalysis (Westerhausen & Papadatou-Pastou, 2022). Irrespective of whether handedness groups were defined based on consistency or direction of hand preference, we did not find statistically significant differences between the compared groups in any of the analyses. Thus, the meta-analysis’ original conclusion that no substantial handedness-related callosal differences can be found, is neither challenged when considering brain size (question 1) nor when conducting regional thickness analyses (question 2). In the following, these findings are discussed in the context of previous findings, the precision of the present analyses, and with respect to theoretical implications.

### 4.1 Accounting for brain-size variability (question 1)

Corpus callosum area and brain size are positive correlated ((e.g., Jäncke et al., 1997; Witelson, 1989), see also Supplementary analysis C) so that considering brain size when comparing the corpus callosum between groups appears intuitively desirable as it might potentially affect the outcome. As discussed by Smith (2005) the inclusion of brain size into the analysis, however, can serve two different purposes: The statistical control of potentially confounding effects by brain-size differences or the analysis of proportionality to brain size. Both approaches will be discussed here in turn.

Statistical control aims to remove the potentially confounding effect of brain size from the analysis, as brain-size differences between handedness groups might increase or decrease group differences in the corpus callosum. It is usually exerted by including brain size measures as covariate in the statistical analysis or by conducting pre-processing step that remove variance attributable to brain-size difference from the dependent measure (e.g., forming residuals, or computational adjustment of the native-space MR images prior to extraction of the corpus callosum, see e.g., (Bermudez & Zatorre, 2001)). Witelson (1985, 1989), introduced brain weight as covariate in the statistical analyses, which, however, did not change the outcome in the statistical decision, as the corpus callosum was still found to be significantly larger in MH compared with cRH. Habib et al. (1991) removed a possible confounding effect by magnifying the MR images so that the midsagittal brain surface area was roughly the same size before measuring the corpus callosum. The authors report the corpus callosum in NcRH to be larger than in cRH, yielding a comparatively large effect size of *d* = 0.88. In the present analyses, comparable to Witelson, we included FBV as covariate to control for brain-size differences and did not find any significant effect of handedness. Hence, we were not able to replicate the previous findings when comparing cRH with MH controlling for brain size. Effect-size estimates reflect small difference, ranging between *d* = 0.04 (equivalent to 3 mm^2^ of absolute mean difference) and 0.07 (6 mm^2^) both in favour of the cRH sample. Also, we did not observe a notable difference to the analysis of absolute callosal area (see Table 1). As mentioned above, Habib et al. (1991) originally compared the cRH with NcRH rather than a MH group, as the author included five cLH participants. To check whether this approach yields different results, we conducted a supplementary analysis (Supplement Section D, Table S4) comparing cRH with NcRH in the present sample. The main effect of Handedness Group was non-significant and the effect size was comparable to the cRH vs. MH findings. Finally, across all analyses, the narrow confidence intervals may be taken to exclude population effects in favour of MH/NcRH above *d* = 0.09 and in favour of cRH above *d* = 0.20. Mean differences that fall in between these values cannot be excluded with confidence, and represent a difference in absolute area of 7 and 17 mm^2^, respectively. Considering a total corpus callosum area of ca. 690 mm^2^ in the present study, this is equivalent to a maximum of 1 to 2% difference in size.

None of the previous studies reporting direction-based handedness group comparisons of the corpus callosum (Denenberg et al., 1991; Jäncke et al., 1997; Mitchell et al., 2003; Moffat et al., 1998; Morton & Rafto, 2006; Nasrallah et al., 1986; Ozdikici, 2020; Tuncer, Hatipoglu, & Ozates, 2005) included brain size as covariate into their analyses. Conducting this analysis for the first time, we here neither for Comparisons C nor D found significant differences, whereby the found empirical effect sizes can be considered small, ranging between *d* = 0.07 (6 mm^2^) and 0.08 (7 mm^2^) in favour of the dRH/cRH as opposed to the dLH/cLH sample. However, the sample size of dLH (n = 106) and cLH (n = 36), were small, resulting in a reduced precision of the analysis reflected by wide confidence intervals of the effect-size estimates. Especially in Comparison D, population differences of up to *d* = 0.27 (24 mm^2^) favouring cLH and up to *d* = 0.41 (35 mm^2^) favouring cRH, cannot be excluded with reasonable certainty. These potential differences represent up to 5% of callosal size.

Rather than being interested in removing the influence of brain size from the analysis, some authors advocate for studying the proportionality of callosal to brain size, emphasising that brain size is relevant for the interpretation of corpus callosum size (Danielsen et al., 2020; Smith, 2005). That is, finding a corpus callosum of identical absolute size in a “small” and a “large” brain would indicate that the small brain has a stronger relative connectivity between hemispheres than the large brain (Aboitiz, 1998; Smith, 2005). Determined by forming ratios of callosal and various brain measures (i.e., FBV, total brain volume, cerebral area on the midsagittal plane) four studies have reported relative callosal area in the context of handedness (Hines, Chiu, McAdams, Bentler, & Lipcamon, 1992; Jäncke et al., 1997; Mitchell et al., 2003; Nasrallah et al., 1986). While some of these report regional differences, none reports significant difference for relative size of the total corpus callosum. Neither Nasrallah et al. (1986) and Mitchell et al. (2003) comparing dRH and dLH, nor Jancke et al. (1997) comparing cRH, MH, and cLH, report significant differences. Also, in the present analyses we do not find significant differences when using ratios of total corpus callosum area to FBV, irrespective of whether a consistency- or a direction-based handedness group comparison was tested. For all four comparisons, the effect sizes were equal to or below *d* = 0.08 favouring the right-handed (cRH, dRH) samples. Comparably to what was discussed above, both consistency-based classifications showed narrow confidence intervals for the effect size not reaching above *d* = 0.19 (for cRH larger than MH) rendering more substantial population effects unlikely. The range of possible effects of the direction-based comparisons are wider (see Table 1), indicating larger uncertainty of the estimates.

Taken together, neither consistency- nor direction-based handedness group comparisons, resulted in significant differences in callosal area when controlling for brain size or when analysing ratios. While the point estimates of effect sizes appear rather small (below *d* = 0.1), the confidence interval (or precision) of the present analyses also deserves consideration as it indicates the range of population effect that cannot be excluded with confidence (Cumming, 2014). For the two consistency comparisons, the confidence intervals allow the exclusion of population effects above *d* = 0.2 in favour of the cRH sample with reasonable certainty. Direction-based comparisons are, however, less precise and the confidence intervals for population effects reach up to *d* = 0.4 favouring cRH/dRH samples. Whether these potential differences can be considered negligible, for example when considering cognitive differences between handedness groups (e.g., Prichard & Christman, 2017; Prichard et al., 2013; Zapała et al., 2020), cannot be answered unequivocally as models linking callosal morphology to cognition or behaviour are missing (Banich, 1998). As proposed by Smith (2005) even small effect sizes may be relevant when accumulated across many events, and inter-hemispheric communication via the corpus callosum might represent such as situation. Nevertheless, compared with other variables known to be associated with callosal size – such as sex (see discussion in section 4.4), brain size (see section 2.6), or aging – the effect sizes are small even considering the extreme limits of the confidence intervals. For example, an effect size of *d* = 0.75 has been reported comparing older (60 years and above) with young (under 30 years) or middle aged (30-60 years) individuals in relative callosal area (Skumlien et al., 2018). Thus, handedness-related callosal variability will likely (if at all) be less relevant for cognition than variability that can be attributed to aging-related atrophy, brain size, or sex differences.

### 4.2 Regional callosal thickness analysis (question 2)

The corpus callosum is topographically organised along its anterior-posterior axes and callosal subsections can be assigned to specific functional brain networks (Schmahmann & Pandya, 2006). Thus, one might predict that especially callosal subregions which link hand-motor cortices (e.g., the truncus of the corpus callosum; see e.g. (Johansen-Berg, Della-Maggiore, Behrens, Smith, & Paus, 2007) are related to handedness rather than the total corpus callosum. To account for this possibility, several studies examined subregional effects by dividing the midsagittal corpus callosum into subsections for a regional analysis. For example, Witelson (1989) conceived a geometrical parcellation schema by dividing the corpus callosum into half, thirds, and fifth relative to the anterior-poster length of its midsagittal surface. Others have applied a principle-component analysis (PCA) to define subregions based on their covariation in callosal thickness (e.g., Cowell & Gurd, 2018; Denenberg et al., 1991).

Regional differences in the size of the handedness effect have accordingly been reported (Habib et al., 1991; Witelson, 1989), and sometimes regional effects without global differences have been found (e.g., Cowell & Gurd, 2018; Hines et al., 1992; Hopper, Patel, Cann, Wilcox, & Schaeffer, 1994; Jäncke et al., 1997; Martens et al., 2013; Moffat et al., 1998). While none of these regional findings have been substantiated in a recent metaanalysis (Westerhausen & Papadatou-Pastou, 2022), it may be argued that the applied geometric rules or PCA-based parcellation do not necessarily reflect functional channels within in the corpus callosum, and that a more fine-grained analysis using thickness measures along the entire corpus callosum offers a spatially more sensitive approach (Luders, Narr, Zaidel, Thompson, Jancke, et al., 2006; Westerhausen et al., 2018). However, only one previous study has applied regional thickness analysis to study effects of handedness. Luders et al. (2010) identified areas in anterior truncus of the corpus callosum in which callosal thickness and the consistency of hand preference (using |LQ| as predictor) were negatively correlated. That is, greater callosal thickness was associated with less consistent hand preference.

Comparing groups of consistent hand preference (comparisons A and B), we here did not find any significant differences in corpus callosum thickness after FDR correction, with or without considering brain size. The effect-size estimates across the 100 segments and both comparisons were below |d| = 0.20. Thus, in contrast to several previous studies comparing cRH with MH or NcRH samples (Habib et al., 1991; Hines et al., 1992; Hopper et al., 1994; Jäncke et al., 1997; Witelson, 1985, 1989), we did not find subregional callosal differences related to handedness. For example, considering absolute thickness, a maximum effect size of *d* = 0.18 (comparison B; see Fig. 2) in favour of cRH over MH was found in the genu which represents a difference of 0.33 mm in a segment with an overall thickness of 10.9 mm. Confidence intervals are in general narrow but extend to an effect of *d* = 0.31 in favour of cRH over MH for the segment showing maximum difference. Of note, Luders et al. (2010) used |LQ| as predictor (instead of or in addition to forming groups) and found regional effects. To allow for direct comparison, we here replicated this approach in a supplementary analysis (Supplement Section E, Fig. S3). Also this analysis failed to detect a substantial association between the degree of hand preference and callosal thickness.

Considering the direction of hand preference, previous studies have found significant regional differences between dRH and dLH samples in truncus and isthmus (Denenberg et al., 1991; Martens et al., 2013; Moffat et al., 1998). However, the direction of the reported difference in the isthmus pointed in opposite directions (compare: Denenberg et al., 1991; Moffat et al., 1998). The present study (comparison C) does not statistically confirm these differences in either direction. Only when considering uncorrected *p*-values, we found a cluster of subthreshold segments in the dorsal splenium, adjacent to the isthmus, being thicker in the dRH than dLH sample. Here the maximum effect size was *d* = 0.26 and the upper confidence limit stretched to *d* = 0.47. For all other callosal subregions the empirical effect sizes are smaller and narrow confidence intervals suggest that populations effects above *d* = 0.30 in favour of dRH may be excluded with confidence.

Finally, Jäncke et al. (1997) found a larger truncus in cRH compared with cLH sample (using a classification comparable to Witelson’s qualitative approach) using both absolute and relative subregional area as dependent measures. Other studies, however, failed to find such effects (Luders et al., 2003; Steinmetz et al., 1992), and in the present study we also did not yield significant difference between cRH and cLH individuals (comparison D) in any of the segments. As discussed for the analysis of total area, the small sample of cLH reduces the precision of the effect size, so that only somewhat large population effects (above *d* = 0.69 in the extreme segment) can be excluded with confidence.

In summary, the present study does not confirm effects of handedness on subregional level, neither when using absolute measures nor when accounting for brain size. The point estimates of the effect sizes across the corpus callosum stay below *d* = 0.2 for consistency-based handedness group comparisons, and the confidence intervals do not extend above *d* = 0.3. Considering comparisons based on the direction of hand preference, empirical effects sizes are slightly higher and associated with lower precision, so that for some thickness segments effects up to *d* = 0.7 cannot be excluded. Thus, while the effect sizes of the consistency-based analyses appear small compared with effects reported for sex (cf. Fig. 4, and discussion in section 4.4) and might be considered less relevant, this is not the case for the direction-based analyses. Thus, although not finding any significant differences, it requires future studies with larger samples of cLH/dLH individuals to narrow down the range of possible effects of the directional hand preference on regional corpus callosum differences.

### 4.3 Defining consistency of hand preference and its callosal effects

Consistency of hand preferences in the past has been either defined based on a qualitative analysis of the participants answer pattern in a handedness questionnaire (e.g., Clarke & Zaidel, 1994; Jäncke et al., 1997; Witelson, 1989) or by using a quantitative definition applying a specific (high) LQ value as cut-off (e.g., Habib et al., 1991; McDowell et al., 2016; Welcome et al., 2009). As we have demonstrated here, these two approaches are not interchangeable, as they may lead to a divergent classification in a substantial number of participants (> 25% in the present implementation, see also Supplement Section B). Previous meta-analyses and reviews on callosal differences in handedness do not distinguish between the two approaches (Budisavljevic et al., 2021; Driesen & Raz, 1995; Westerhausen & Papadatou-Pastou, 2022) so that potential differences between qualitative and quantitative classifications could have been overlooked. However, across all analyses presented here, we do not find any indication for this, as the findings of Comparison A and B appear highly comparable. For example, regarding the analysis of absolute area using the qualitative definition the effect size was *d* = 0.04 while it was *d* = 0.06 for the quantitative definition, with largely overlapping confidence intervals, and both favouring the cRH over the MH sample (see Table 1).

In context of quantitative definitions, it also deserves discussion that the setting of the LQ cut-off value is in the end arbitrary, and various LQ thresholds have been used in the literature (Hardie & Wright, 2014). In the present paper, by using an LQ of 80 and above to define consistent hand preference, we followed the definition by Habib et al. (1991), who do not provide an explicit justification for the choice of the threshold. However, both more inclusive and more restrictive definitions of consistency can be found, and threshold values between 75 (Propper & Christman, 2004) and 100 (McDowell et al., 2016; Welcome et al., 2009) have been selected. Thus, we were curious whether choosing a different cut-off value might alter the outcome. To test a wide range of possible values, we conducted an explorative analysis applying |LQ|-thresholds between 10 and 100 separating groups of “consistent” from groups of non-consistent hand preference. As outlined in Supplementary Section F (e.g., Fig. S4, we did not find significant differences in corpus callosum size for any of the applied thresholds.

### 4.4 Moderating and main effect of Sex

A series of early studies suggest that the effect of handedness on the total corpus callosum size, but especially on the size of the isthmus subregion, is moderated by the participants’ sex (Burke & Yeo, 1994; Clarke & Zaidel, 1994; Denenberg et al., 1991; Habib et al., 1991; Witelson, 1989; Witelson & Goldsmith, 1991). Across these studies it appeared that handedness differences in callosal size were stronger in males than females or pointed in opposite directions for the two sexes. For example, Witelson (1989) reported the isthmus in male but not in female brains tended to be larger in MH than in cRH. Burke and Yeo (1994) found a positive correlation of hand preference (LQ as continuous variable) and callosal area in their male subsample, while a reversed association was found in the female sample. In the present study we do not find any indication that Sex might act as moderator variable, as the interaction of Handedness and Sex was not significant in any of the analyses and the corresponding effect size estimate were well below 1% explained variance across analyses (see Table 1). Thus, neither considering total callosal area, nor when selectively considering the thickness segments representing this isthmus, were we able to replicate the findings of the above studies.

Although not finding an interaction, we did find a main effect of Sex. Regarding absolute midsagittal area, the corpus callosum was larger in male than female participants, but the effect was found reversed when considering brain size (as covariate or when using relative size). This pattern is well in line with previous meta-analyses on callosal sex differences. That is, absolute area is typically found to be larger in males, and analyses of relative area and using brain size as covariate tend to find larger areas in females (Driesen & Raz, 1995; Bishop & Wahlsten, 1997; Smith, 2005). Furthermore, the present effect sizes of the Sex effect were in the same range as the estimates of these earlier meta-analyses. That is, the *d* = 0.23 found for absolute area, is comparable the effect size estimate of the largest meta-analysis on absolute measures by Bishop & Wahlsten (1997) which found a *d* = 0.21 (*CI_95%_*: 0.13; 0.29) in favour of males. The effect sizes for the analyses considering brain size, were with *d* = 0.49 (covariate FBV) and 0.54 (relative area) in favour of females comparable to the findings by Smith (2005), who estimated *d* of the sex difference between 0.22 (*CI_95%_*: 0.07; 0.40) and 0.62 (*CI_95%_*: 0.47; 0.75) in his series of meta-analyses comparing brain size adjustment methods. Notably, however, the sample sizes included in the meta-analyses by Smith (2005) are of same size or even smaller than in the present sample, so that the present estimates might be seen more informative than the meta-analyses.

Previous studies using callosal thickness measures reflect this general pattern of the area analyses, as absolute thickness (across most of the corpus callosum) is typically found larger in males (Luders, Narr, Zaidel, Thompson, & Toga, 2006; Westerhausen et al., 2016), while once brain size is corrected, the sex effects vanishes (Luders, Toga, & Thompson, 2014; Westerhausen et al., 2016) or is reversed (Danielsen et al., 2020; Luders, Narr, Zaidel, Thompson, & Toga, 2006). In the present study, only segments in the splenium were significantly thicker in males than females when analysing absolute thickness, but several segments in the genu showed empirical effects of *d* = 0.3 and above but did not survive FDR adjustment. Introducing FBV as covariate or analysing relative thickness, reversed the effect, indicating significantly thicker female corpora callosa in most segments, from genu to splenium. The effect sizes of significant difference were larger than *d* = 0.3, reaching up to *d* = 0.5 in favour of females.

In summary, while there is no indication for differential effects of handedness in the two sexes, the main effect of Sex replicate previous findings both when using absolute measures and when accounting for brain size differences. The corpus callosum tends to be larger or thicker in males when brain size is ignored, while it is larger in females when brain size is accounted for. The relatively stronger structural connectivity between the hemispheres in the female brain, has been previously attributed to reflect difference in brain size rather than being a sex-specific effect (Hänggi, Fövenyi, Liem, Meyer, & Jäncke, 2014; Jäncke et al., 2015; Jäncke et al., 1997; Luders, Narr, Zaidel, Thompson, & Toga, 2006). In an influential paper, Ringo, Doty, Demeter, and Simard (1994) proposed that maintaining interhemispheric connectivity in larger compared to smaller brains becomes computationally inefficient, as the increased inter-cortical distances increase the conduction delays to a degree that they cannot be compensated. As consequence, larger compared with smaller brains exhibit reduced inter- and enhanced intra-hemispheric connectivity (Hänggi et al., 2014) and the ratio of callosal to brain size is accordingly decreased in larger brains (Jäncke et al., 1997; Leonard et al., 2008). As the female brain is on average smaller than the male (Jäncke et al., 2015), the here found sex differences in the corpus callosum in favour of females, potentially reflects the higher ratio of callosal size to brain volume in smaller brains.

### 4.5 Limitations

As outlined in the introduction, defining handedness based on direction of hand preference does not provide a straight-forward prediction regarding callosal morphology without making additional assumptions concerning other functional brain modules. For example, assuming that the language planning is controlled dominantly by the left hemisphere, dLH compared with dRH individuals would require additional communication – and stronger callosal connectivity – between the hemispheres when writing with the left hand (controlled by the right hemisphere). However, dLH and dRH individuals differ in their brain organisation also regrading other modules, such as language processing, as a substantial larger amount of dLH (ca. 30-35%) than dRH (ca. 5-10%) are not left-lateralised for language (Carey & Johnstone, 2014). Given the writing example, only dLH with left hemisphere language and dRH with right hemisphere language processing would have to rely on callosal transfer. Thus, in the present analyses by only considering handedness, we formed heterogenous groups in which callosal effect would not be predicted for all members, potentially masking existing association. Consequently, future studies of the corpus callosum could benefit from analysing the effect of handedness in interaction with the lateralisation of other functional brain modules (Gerrits et al., 2020; Karlsson et al., 2022).

One limitation of the present study is that handedness was only operationalized by hand preference questionnaires, ignoring that hand preference does not perfectly reflect dexterity differences between the hands (Andersen & Siebner, 2018). Alternative measures have been conceptualized assessing left-right hand-skill differences for certain motor tasks (such as pegboard and drawing tasks; (e.g., Annett, 1976; McManus, Van Horn, & Bryden, 2016)), and previous studies have related hand-skill differences to callosal morphology, although without finding any substantial association (Kertesz et al., 1987; Preuss et al., 2002; Steinmetz, Staiger, Schlaug, Huang, & Jäncke, 1995). Nevertheless, as the present aim was to comprehensively assess the relation of handedness and the corpus callosum, omitting skill-based handedness can be seen as a short-coming of the present study, especially since performance and self-report handedness not necessarily correlate well (Corey, Hurley, & Foundas, 2001). While the HCP database provides measures that may potentially be used to determine hand-skill differences (a pegboard task and measures of grip strength from the NIH toolbox, (Gershon et al., 2010)), a recent analysis of the HCP data demonstrated low retest reliability for laterality measures in the two tasks. That is, a Pearson correlations of .28 and .42 for the pegboard task and grip strength, respectively, were reported (Ruck & Schoenemann, 2021). Given this low reliability, we refrained including skill-based handedness classifications in the present analyses. Thus, the present study was unfortunately not able to provide more evidence on the association of skill laterality and the corpus callosum.

## 5 Conclusion

Overall, our findings support the conclusion of the meta-analysis that motivated the present study (Westerhausen & Papadatou-Pastou, 2022) as we were not able to detect handedness-related callosal differences. That is, neither when considering brain-size difference in our analysis nor when analysing spatial-sensitive regional thickness measures, did we detect significant group differences. As discussed, the interpretation of these null findings also hinges on the statistical precision, that is, the range of population effect sizes than can be included with reasonable certainty (Cumming, 2014). For consistency-based definitions of hand preference these excludable population effects appear, in our opinion, reasonably small to suggest that they are negligible for explaining difference in behaviour and cognition. For direction-based definitions confidence intervals are wider as the precision is reduced due to small number of cLH/dLH individuals in the sample. Thus, future studies including large samples of left handers (cLH/dLH)) might help to further narrow down the possible population effects. In any case, it can be concluded that the common practise of inferring callosal connectivity from handedness (Jasper et al., 2021; Roberts et al., 2020; Zapała et al., 2020) is not justified and should be avoided as long as no actual measures of the corpus callosum are analysed.

## Supporting information

Suppl analyses

## 6 Acknowledgment

The data was kindly provided by the Human Connectome Project, WU-Minn Consortium (Principal Investigators: David Van Essen and Kamil Ugurbil; 1U54MH091657) funded by the 16 NIH Institutes and Centers that support the NIH Blueprint for Neuroscience Research; and by the McDonnell Center for Systems Neuroscience at Washington University.

