## Supplementary material for "Hand preference and the corpus callosum: Is there really no association?": Suppl analyses

#### A. HCP handedness questionnaire and determination of the laterality quotient (LQ) for the present analyses

*Background*. The variable "Handedness" provided by the Human Connectome Project (HCP) according to the Data Release Documentation represents the laterality quotient (LQ) of the Edinburgh Handedness Inventory (EHI, Oldfield, 1971). However, raw item-wise data for only nine of the ten manual preference items are available, leaving out the item "drawing" from the original item set (see also Ruck & Schoenemann, 2021). Thus, given the missing item, it was not immediately clear how the LQ was formed. In interaction with the HCP team, and as could be verified by own calculations, it turned out that the HCP LQ is calculated by including the "kicking with foot" item in addition to the 9 "manual" items. Thus, the LQ provided by HCP as "Handedness" is a mixture of manual and pedal preference.

As footedness and handedness are not perfectly related (Packheiser et al., 2020), it was necessary to determine a new LQ based on the nine available manual items for the present study. To this end, we explored how the data of the available nine manual items in HCP can be used to obtain a predictor comparable to the full-scale LQ. To this end, an unpublished data set collected with a Norwegian version of the 10-item EHI was utilized to calculate the correlation of the answers of the ten items. Spearman rank correlations were used to account for the non-normal distribution of the EHI item scores. The correlation matrix in Table S1 shows that the answers on the "drawing" item correlate strongly with the answers to the item "writing" ($r_{sp}$ =.89), but less strongly with the other eight items ($r_{sp}$ between .29 and .59). Also, only four of the 97 (4.4%) participants scored differently on the items asking for "writing" and "drawing", and none of these four indicated to prefer opposite hands for the two activities. Thus, it appears the "writing" and "drawing" items are somewhat redundant in the EHI.

*Table S1*. Spearman correlation ($r_{sp}$) of answers to the 10 EHI questions in sample of N=97 participants [a]

| | EHI items | | | | | | | | | |
|---|---|---|---|---|---|---|---|---|---|---|
| | wri | dra | thr | sci | too | kni | spo | bro | mat | box |
| writing | 1 | 0.89[b] | 0.50 | 0.45 | 0.59 | 0.53 | 0.55 | 0.42 | 0.54 | 0.29 |
| drawing | | 1 | 0.59 | 0.48 | 0.51 | 0.62 | 0.53 | 0.39 | 0.56 | 0.27 |
| throwing | | | 1 | 0.36 | 0.40 | 0.39 | 0.4 | 0.31 | 0.41 | 0.39 |
| scissors | | | | 1 | 0.36 | 0.41 | 0.41 | 0.39 | 0.42 | 0.31 |
| toothbrush | | | | | 1 | 0.46 | 0.54 | 0.41 | 0.51 | 0.30 |
| knife | | | | | | 1 | 0.52 | 0.31 | 0.59 | 0.20 |
| spoon | | | | | | | 1 | 0.45 | 0.52 | 0.32 |
| broom | | | | | | | | 1 | 0.50 | 0.41 |
| match | | | | | | | | | 1 | 0.29 |
| box | | | | | | | | | | 1 |

*Notes.* (a) unpublished dataset, including data from 67 female and 30 male healthy participants, mean age: 24.8 years (standard deviation: 3.8 years); (b) all correlation p < .05 uncorrected.

From this observation it was decided to determine the new LQ for the HCP data by double weighing the answer of "writing" when calculating the LQ from the nine items available. We argue that this approach biases the resulting LQ less than using the answers to the nine items with an equal weight. That is, including all items with equal weight would bias the LQ towards activities which are less correlated with other manual activities and that have been referred to as "secondary" manual activities (Annett, 1970). Rather, substituting the missing “drawing” item with a second counting of “writing” item should yield an LQ from the 9 items that is comparable to the LQ obtained from the full-scale EHI.

*Approach*: To test this prediction, we used the above data set to calculate the (1) full-scale LQ ($LQ_{standard}$), (2) an LQ based on equal weight of the nine items available from HCP ($LQ_{equal}$), and (3) and LQ based on double weighing the “writing” item ($LQ_{double\_writing}$). All three LQ variables were scaled to range from -100 to 100 for consistent left-hand and right-hand preference, respectively.

*Results*. While both $LQ_{equal}$ and $LQ_{double\_writing}$ correlate highly with the $LQ_{standard}$ (both $r_{sp} > .99$, $p < .001$), paired t-tests indicated that $LQ_{equal}$ differed significantly from $LQ_{standard}$ ($t_{96} = 6.28$, $p < .001$), with higher values for the standard LQ (difference: 1.44; $CI_{95\%}$ of difference: 0.98, 1.89). However, $LQ_{double\_writing}$ did not differ significantly from $LQ_{standard}$ ($t_{96} = 1$, $p = .32$) whereby the standard LQ was slightly larger (difference: 0.10; *CI95%*: -0.10, 0.31).

*Conclusion*. Although both approaches to calculate an LQ based on the available nine items yield LQs that are highly correlated with the standard (full-scale) LQ, the existence of mean differences between $LQ_{equal}$ and $LQ_{standard}$ might lead to differential classification of participants when thresholds are used. The $LQ_{double\_writing}$, showing a small (non-significant) mean deviation from $LQ_{standard}$, renders differences in classification less likely, and was accordingly used in the present study.

### B. Inter-relation between various methods of classifying hand preference

*Consistency-based classification* of hand preference groups may either be based on a qualitative analysis of the individual answer pattern (e.g., Clarke & Zaidel, 1994; Jäncke, Staiger, Schlaug, Huang, & Steinmetz, 1997; Witelson, 1989) or by using a specific LQ value as threshold (e.g., Habib et al., 1991; McDowell, Felton, Vazquez, & Chiarello, 2016; Welcome et al., 2009). Here, as outlined in the main manuscript, we utilized both approaches, and the following Table S2 illustrates that a substantial proportion of participants are classified differently by the two approaches.

*Table S2*. Classification of participants based on qualitative (Witelson) and quantitative (Habib) definition

| | | Habib criterion | | | |
|---|---|---|---|---|---|
| | | cRH | MH | cLH | Sum |
| Witelson | cRH | **502**[a] | 134 | 0 | *636* |
| criterion | MH | 129 | **249** | 7 | *385* |
| | cLH | 0 | 7 | **29** | *36* |
| | Sum | *631* | *390* | *36* | 1057 |

*Notes*. cRH: consistent right handers, cLH: consistent left handers, MH: "mixed" handers; a: bold face numbers: same classification according to both criteria

While leading to very similar marginal frequency distributions, the off-diagonal elements show that the classification differ between both criteria. That is, 277 out of 1057 participants (26.2%) were classified differently by the two criteria. As this difference cannot be ignored, separate statistical analyses for both criteria appear warranted.

More recent studies rather than assessing consistency of hand preferences, aimed to compare groups based on the *direction of preference*. That is, groups were defined by the preferred hand for writing (e.g., Cowell & Gurd, 2018; Denenberg, Kertesz, & Cowell, 1991) or the overall preference across multiple activities (e.g., as expressed by the sign of the LQ) (e.g., Martens, Wilson, Chen, Wood, & Reutens, 2013; Moffat, Hampson, & Lee, 1998; Nasrallah et al., 1986). A comparison of both approaches can be seen in Table S3.

*Table S3*. Classification of participants based on writing hand or the sign of the LQ

| | | Sign of LQ | | | |
|---|---|---|---|---|---|
| | | dRH | np | dLH | Sum |
| Writing | dRH | **930**[a] | 0 | 3 | *933* |
| hand | np | 1 | **1** | 0 | *2* |
| | dLH | 15 | 4 | **103** | *122* |
| | Sum | *946* | *5* | *106* | 1057 |

*Notes*. dRH: dominant right handed, dLH: dominant left handed, np: no preference; a: bold face numbers: same classification according to both criteria

As can be seen 23 participants were classified differently by the two approaches, that is equivalent to 2% of the cases. Thus, writing hand and sign criterion produce largely overlapping classifications, so that we only utilized the classification based on the sign of the EHI, as a classification on multiple activities appears more reliable than just focussing on writing.

### C. Forebrain volume as predictor of corpus callosum area and thickness

To illustrate the effect of forebrain volume (FBV), we conducted simple linear regression analyses predicting total corpus callosum area and regional callosal thickness from FBV. FBV was found to be significant predictor of total callosal area ($b = 0.00016$, $se(b) = 0.000014$, $t_{1055} = 11.11$, $p < .001$) explaining 10.5% of the variance (i.e., $R^2$). As we used adjusted FBV ($FBV^{2/3}$) to correct for brain size in the main analyses, we repeated the analysis using this predictor and yielded a comparable result ($b = 0.046$, $se(b) = 0.004$, $t_{1055} = 12.59$, $p < .001$, $R^2 = .13$). The scatterplots are shown in Figure S1.

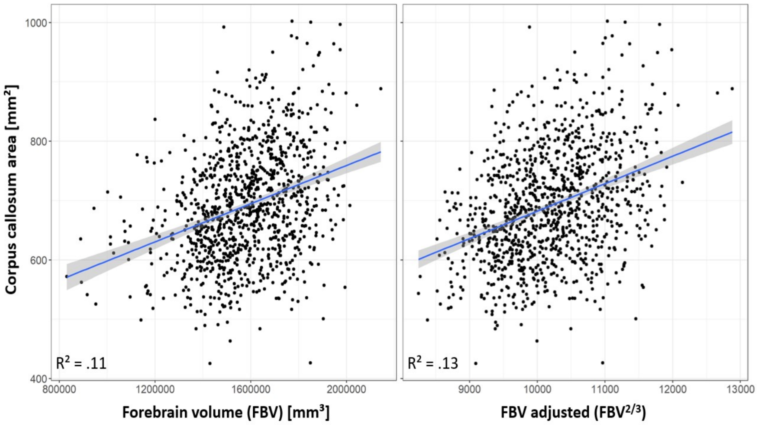


*Fig. S1*. Prediction (blue line) of the total corpus callosum area from FBV, both using absolute and adjusted FBV. The shaded area represents the 95% confidence band of the prediction.

Repeating the same analyses for the thickness measures (see Fig. S2) we found segment thickness throughout genu, truncus, and splenium to be significantly predicted by FBV after FDR adjustment, with a maximum of $R^2 = .09$. Using the adjusted FBV (here $FBV^{1/3}$) comparable results were found, again explaining up to 9% variance in callosal thickness.

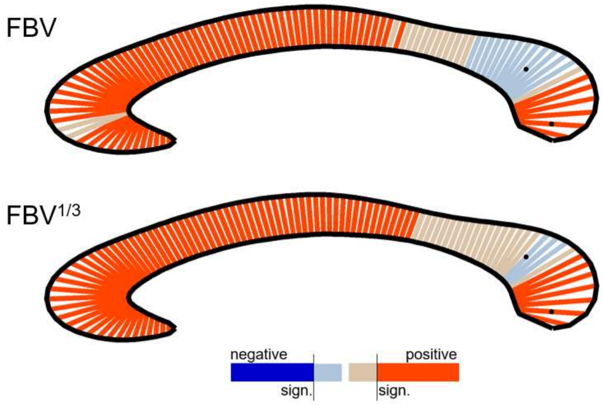


*Fig. S2*. Regional thickness predicted by FBV, using absolute and adjusted FBV as predictors. Significance level adjusted to an FDR of 5%. Black dots mark the segment showing the largest effect in either direction. The $R^2$ for significant associations ranged from .004 to .09.

**D. Supplementary analysis: Conceptual replication of Habib et al. (1991)**

*Background*: Habib et al. (1991) did compare cRH with NcRH individuals, that is, the authors did not distinguish between cLH and MH. However, being interested in the effect of "consistency" the comparison with an NcRH sample appears not optimal as both individual with inconsistent (MH) and consistent (cLH) hand preference are included in the same group. Consequently, in the present analysis we decided to set up Comparison B of the main analysis as cRH vs. MH instead. For reasons of comparability with the original publication, we here nevertheless provide the cRH vs. NcRH comparison, which may be seen as conceptual replication attempt. Of note, as Habib et al. (1991) accounted for brain-size differences by normalizing the brain images to the same midsagittal size before measuring the corpus callosum. This approach appears may be seen as correction for brain size by removing its variance from the data which is best compared with the analysis using FBV estimates as covariate.

*Approach*: As for the main analyses discussed in the manuscript, analyses of variance (ANOVA) with the factors Handedness Group (here levels: cRH and NcRH) and Sex were used for analysis, while using absolute and relative area of the corpus callosum as dependent variable in separate analyses. A third analysis was conducted using absolute area as dependent variable but adding the converted FBV as covariate into the design. Across all three analyses, the effect of interest was the main effect of Handedness Group.

*Results*: For neither of the analyses a significant main effect of Handedness Group or interaction with Sex was found as indicated in the following table:

*Table S4*. Results comparing cRH and NcRH following the approach by Habib et al. (1991)

| | Main effect Handedness Group (HG) | | | | | Interaction HG x Sex | |
|---|---|---|---|---|---|---|---|
| Analysis | *F* value | $df_n$, $df_d$ | *p* | *d* | *CI95%(d)* | *p* | $\eta^2$ |
| Absolute area | 1.44 | 1; 1053 | 0.23 | 0.08 | -0.05; 0.20 | 0.91 | <0.001 |
| Relative area | 1.25 | 1; 1053 | 0.26 | 0.07 | -0.05; 0.20 | 0.73 | <0.001 |
| $FBV^{2/3}$ as covariate | 1.51 | 1; 1052 | 0.22 | 0.08 | -0.05; 0.20 | 0.68 | <0.001 |

*Notes*. The main effect of sex was significant for all analyses, following the same pattern as reported in the main manuscript. Cohen's *d* effect size is calculated based on the estimated marginal means with positive values indicating larger areas in cRH. Please see accompanying script/R markdown document for mean values and more details.

*Conclusion*: Habib et al. (1991) reported an effect size of *d* = -0.87, indicating lager total corpus callosum area for NcRH compared with cRH. Here, we were not able to replicate the Habib et al. (1991) findings. That is, across all three dependent variables, the direction of the effect is reversed in showing larger area measures in cRH than NcRH, and the confidence interval in fact exclude effects in favor of NcRH stronger than *d* = -0.05.

**E. Supplementary analysis of thickness measures: conceptual replication of Luders et al. (2010)**

*Background*: Luders et al., (2010) reported negative correlations between absolute EHI score (i.e., a marker of consistency of hand preference ignoring the direction) and thickness in several segments in the truncus of the corpus callosum. The sample consisted of N = 361 participants with EHI LQ varying between -100 to 100, and the authors indicate that 324 participants were right and 37 left handers (by direction of preference). The statistical analysis was described as: *"… we converted the directional EHI-handedness measures into absolute values … Then, we calculated the correlation coefficients between the absolute EHI scores and callosal thickness at 100 surface points within the overall sample …, while covarying for ICV* [intra-cranial volume]" (p. 45).

*Approach*: We here replicated this approach by setting up segment-wise linear regressions with the absolute value of the LQ (i.e., |LQ|). To account for brain-size effects, we entered estimated ICV (HCP variable: *FS_IntraCranial_Vol*) as additional predictor into the model. Thus, in contrast to all other analyses in which FBV was used as estimate of brain size, we here used ICV to closely follow the procedure described by Luders et al.

*Results*: For none of the segments a significant effect of |LQ| was found after applying FDR correction (to 5%) for multiple comparisons. Considering uncorrected *p*-values for two segments located in the splenium of the corpus callosum (maximum located at first dotted segment in Fig. S3), a positive association of |LQ| and thickness was significant, while one segment in the ventral splenium showed a negative association (second dotted segment). However, the maximal effect size (found at segment #99) was $\eta^2 = 0.007$, that is, explaining less than 1% variance in the data.

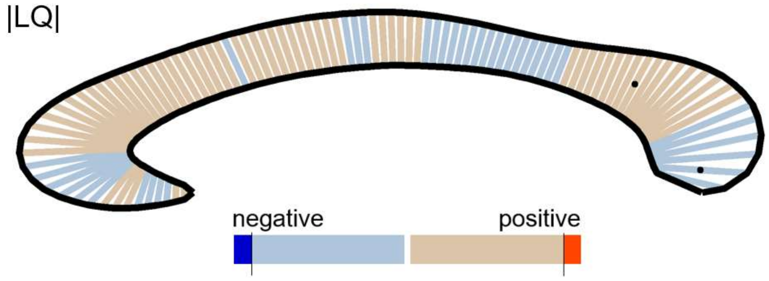


*Fig. S3*. Regional thickness predicted by |LQ|. Significance level adjusted to an FDR of 5%. Black dots mark the segment showing the biggest effect in either direction.

*Conclusion*: The present attempt of replication did not confirm the findings reported by Luders et al. (2010).

**F. Supplementary analysis: effect of LQ threshold on consistency- and direction-based analyses**

*Consistency of hand preferences*

Considering the quantitative approach, the LQ-threshold used to separate cRH (or cLH) from mixed handed groups is set arbitrarily and accordingly various thresholds have been used in the past (Habib et al., 1991; Propper & Christman, 2004; Welcome et al., 2009). To examine the effect the threshold has on possible callosal differences, we here conduct an explorative analysis using LQ-thresholds between 10 and 100 to separate individuals of consistent hand preference (cH) from non-cH (or NcH) preference. Individuals with an |LQ| equal or larger to the threshold were classified as cH, and all others as NcH. Joining cLH and cRH into one group emphasizes consistency of hand preference over direction and has been employed in previous studies (McDowell et al., 2016; Welcome et al., 2009). The numbers of participants per group as function of the threshold can be seen in Table S5 .

*Table S5*. Number of cH and NcH participants by threshold

| | \|LQ\| threshold | | | | | | | | | |
|---|---|---|---|---|---|---|---|---|---|---|
| | 10 | 20 | 30 | 40 | 50 | 60 | 70 | 80 | 90 | 100 |
| cH | 1044 | 1035 | 1021 | 1003 | 970 | 922 | 836 | 667 | 427 | 249 |
| NcH | 13 | 22 | 36 | 54 | 87 | 135 | 221 | 390 | 630 | 808 |

*Notes*. cH = consistent hand preference, NcH = non-consistent hand preference.

Two-factorial analyses of variance (ANOVAs) with the factor Handedness Group and Sex were run for each resulting classification and for the two dependent variables absolute and relative area (of the total corpus callosum). Adding $FBV^{2/3}$ as covariate to the design, the analysis of absolute area was repeated as analysis of covariance (ANCOVA). Figure S4 shows the Cohen's *d* for the Handedness Group comparisons, as calculated from the estimated marginal means for cH and NcH. As reflected by the 95% confidence limits always including "0", no significant (all uncorrected $p > .05$ for the main effect) Handedness Group difference could be found.

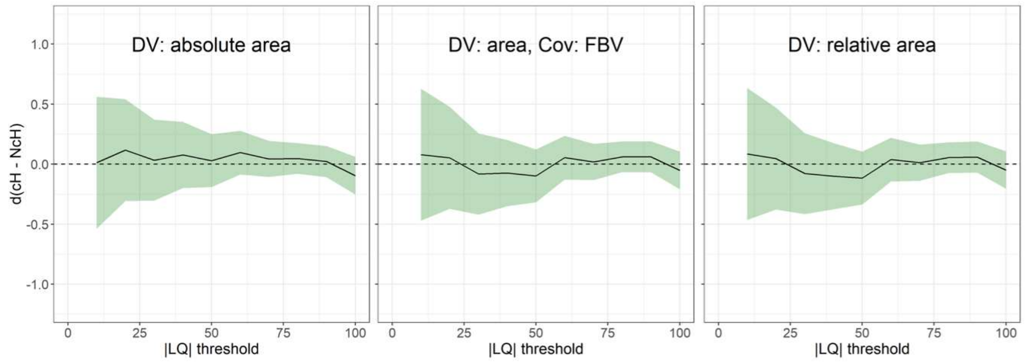


*Fig. S4*. Cohen's *d* for the comparison of cH and NcH samples as function of the |LQ| threshold used to define the two groups (black line). Green-shaded areas mark the 95% confidence of *d*. Positive values of *d* indicate larger area in cH than NcH while negative values indicate the reversed difference. The results are presented separately for the three analyses for the two dependent variables (DV) and when using FBV as covariate (Cov).

Across all three analyses the confidence intervals are wide for lower thresholds, which can be attributed the comparatively small NcH samples below the |LQ| = 50. From |LQ| of 60 and above, however, whin the range of *d* =-0.25 and 0.25.

In conclusion, irrespective of which LQ threshold is used to defined consistent hand preference, the empirical effect in the present sample is small. For LQ thresholds typically used in the literature to define consistency substantial population-level difference between cH and NcH appear unlikely.

*Direction of hand preferences*

Following the above approach, we also evaluated the effect the LQ threshold has for the comparison of groups difference in the direction of hand preference (e.g., dominant left vs. dominant right handers, or consistent left vs. consistent right handers). That is, using LQ thresholds mirrored at zero, we defined left handers (LH; i.e., an individual LQ equal and below the negative value of a given LQ threshold) and right handers (RH, i.e., LQ equal and above the positive LQ thresholds), while using threshold values between |LQ| = 0 and |LQ| = 100. Table S6 shows the numbers of participants per group as function of the used threshold.

*Table S6*. Number of RH and LH participants by |LQ| threshold

| | \|LQ\| threshold | | | | | | | | | | |
|---|---|---|---|---|---|---|---|---|---|---|---|
| | 0 | 10 | 20 | 30 | 40 | 50 | 60 | 70 | 80 | 90 | 100 |
| RH | 946 | 944 | 939 | 931 | 921 | 900 | 860 | 782 | 631 | 402 | 234 |
| LH | 106 | 100 | 96 | 90 | 82 | 70 | 62 | 54 | 36 | 25 | 15 |
| Excluded[1] | 5 | 13 | 22 | 36 | 54 | 87 | 135 | 221 | 390 | 630 | 808 |

Notes. (1) Participants below |LQ| threshold, i.e. "mixed handers" were excluded from the analyses. RH = right handers, LH = left handers.

The statistical analysis was done using two-factorial ANOVAs with the factor Handedness Group (RH vs LH, excluding all others) and Sex for each threshold as well as for the two dependent variable (absolute and relative area). The analysis of absolute area was repeated using $FBV^{2/3}$ as covariate making it an ANCOVA. An overview of the results can be seen in Fig. S5.

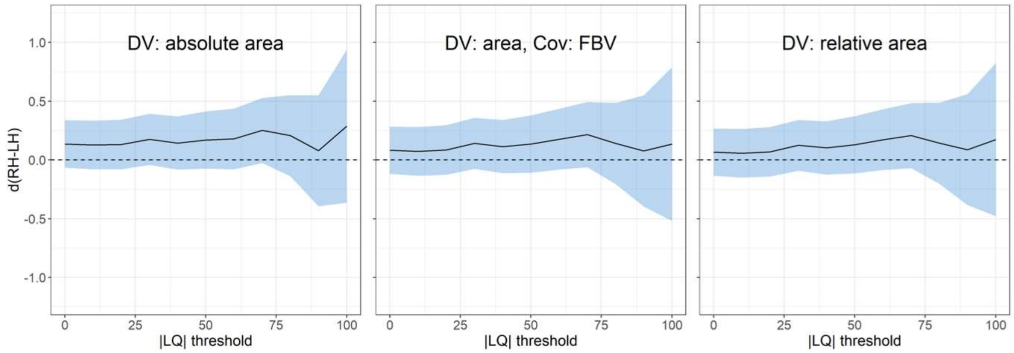


*Fig. S5*. Cohen's *d* for the main effect of Handedness Group, as function of the |LQ| threshold used to define the two groups (black line; shaded areas: 95% CI). Positive *d*s indicate larger area in RH than LH and negative values indicate the reverse. The results are presented separately for the three analyses for the two dependent variables (DV) and when using FBV as covariate (Cov). Of note, the analyses/results for the threshold of LQ = 80 are equivalent to those of Comparison D presented in te main text; the results for the threshold of LQ = 0 are equivalent to those of Comparison C.

Across all analyses the main effect of Handedness Group was non-significant (all uncorrected $p > .05$ for the main effect). As can be seen in Figure S5, the estimated effect sizes were positive (i.e., RH > LH) in all analyses whereby the maximal effect size did not exceed above $d = 0.29$. The confidence intervals for $d$ are wide for thresholds of (roughly from an |LQ| of 60) which can be attributed to a small number of LH participants and an increasing number of exclusions (see Table S6). However, up to and including the threshold of |LQ| = 50, the upper limit does not reach $d = 0.40$, while the lower limit does not reach $d = -0.10$.

In conclusion, all analyses were non-significant and the found effect sizes small. For lower thresholds, the confidence limits further suggests that substantial population effects are unlikely, especially, in favour of LH may be excluded.

### References Supplement


Annett, M. (1970). A classification of hand preference by association analysis. *British journal of psychology, 61*(3), 303-321.

Clarke, J. M., & Zaidel, E. (1994). Anatomical-behavioral relationships: corpus callosum morphometry and hemispheric specialization. *Behav Brain Res, 64*(1-2), 185-202. doi:10.1016/0166-4328(94)90131-7

Cowell, P. E., & Gurd, J. (2018). Handedness and the Corpus Callosum: A Review and Further Analyses of Discordant Twins. *Neuroscience, 388*, 57-68. doi:10.1016/j.neuroscience.2018.06.017

Denenberg, V. H., Kertesz, A., & Cowell, P. E. (1991). A factor analysis of the human's corpus callosum. *Brain Res, 548*(1-2), 126-132. doi:10.1016/0006-8993(91)91113-f

Habib, M., Gayraud, D., Oliva, A., Regis, J., Salamon, G., & Khalil, R. (1991). Effects of handedness and sex on the morphology of the corpus callosum: a study with brain magnetic resonance imaging. *Brain Cogn, 16*(1), 41-61.

Jäncke, L., Staiger, J. F., Schlaug, G., Huang, Y. X., & Steinmetz, H. (1997). The relationship between corpus callosum size and forebrain volume. *Cerebral Cortex, 7*(1), 48-56. doi:10.1093/cercor/7.1.48

Martens, M. A., Wilson, S. J., Chen, J., Wood, A. G., & Reutens, D. C. (2013). Handedness and corpus callosal morphology in Williams syndrome. *Development and Psychopathology, 25*(1), 253-260. doi:10.1017/s0954579412001009

McDowell, A., Felton, A., Vazquez, D., & Chiarello, C. (2016). Neurostructural correlates of consistent and weak handedness. *Laterality, 21*(4-6), 348-370. doi:10.1080/1357650x.2015.1096939

Moffat, S. D., Hampson, E., & Lee, D. H. (1998). Morphology of the planum temporale and corpus callosum in left handers with evidence of left and right hemisphere speech representation. *Brain, 121*, 2369-2379. doi:10.1093/brain/121.12.2369

Nasrallah, H. A., Andreasen, N. C., Coffman, J. A., Olson, S. C., Dunn, V. D., Ehrhardt, J. C., & Chapman, S. M. (1986). A controlled magnetic resonance imaging study of corpus callosum thickness in schizophrenia. *Biol Psychiatry, 21*(3), 274-282. doi:10.1016/0006-3223(86)90048-x

Oldfield, R. C. (1971). The assessment and analysis of handedness: the Edinburgh inventory. *Neuropsychologia, 9*(1), 97-113.

Propper, R., & Christman, S. (2004). Mixed-versus strong right-handedness is associated with biases towards" remember" versus" know" judgements in recognition memory: Role of interhemispheric interaction. *Memory, 12*(6), 707-714.

Welcome, S. E., Chiarello, C., Towler, S., Halderman, L. K., Otto, R., & Leonard, C. M. (2009). Behavioral correlates of corpus callosum size: Anatomical/behavioral relationships vary across sex/handedness groups. *Neuropsychologia, 47*(12), 2427-2435. doi:10.1016/j.neuropsychologia.2009.04.008

Witelson, S. F. (1989). Hand and sex differences in the isthmus and genu of the human corpus callosum. A postmortem morphological study. *Brain, 112 ( Pt 3)*, 799-835. doi:10.1093/brain/112.3.799
